# Studying Spatial Memory in Augmented and Virtual reality

**DOI:** 10.1101/777946

**Authors:** Shachar Maidenbaum, Ansh Patel, Isaiah Garlin, Josh Jacobs

## Abstract

Spatial memory is a crucial part of our lives. Spatial memory research and rehabilitation in humans is typically performed either in real environments, which is challenging practically, or in Virtual Reality (VR), which has limited realism. Here we explored the use of Augmented Reality (AR) for studying spatial cognition. AR combines the best features of real and VR paradigms by allowing subjects to learn spatial information in a flexible fashion while walking through a real-world environment. To compare these methods, we had subjects perform the same spatial memory task in VR and AR settings. Although subjects showed good performance in both, subjects reported that the AR task version was significantly easier, more immersive, and more fun than VR. Importantly, memory performance was significantly better in AR compared to VR. Our findings validate that integrating AR can lead to improved techniques for spatial memory research and suggest their potential for rehabilitation.

**Highlights:**

- We built matching spatial memory tasks in VR and AR
- Subjectively, subjects find the AR easier, more immersive and more fun
- Objectively, subjects are significantly more accurate in AR compared to VR
- Pointing based tasks did not fully show the same advantages
- Only AR walking significantly correlated with SBSoD, suggesting mobile AR better captures more natural spatial performance

## Introduction

Where did I leave my keys? Where did I park my car? As we go about our day we are constantly faced with spatial memory tasks, in which we form associations between various objects with specific locations. To understand how spatial memory works, one can perform experiments in the real world, by placing items in different locations and asking subjects to remember a given object’s location. However such real-world experiments are inherently cumbersome—to test spatial memory for multiple items in different locations one would need to collect those items, manually position them around the environment for each trial, all while being restricted by physical limitations such as the available environment, and equipment. These technical difficulties have led to a popular use of virtual-reality (VR) based paradigms for studying spatial memory (e.g. [16,25,30]). On the one hand, these VR paradigms are much more flexible and easy to run. On the other hand, spatial memory and navigation in VR is inherently different compared to the real world. VR lacks the physical motion, level of immersion, and idiothetic (internal self motion) cues of real-world navigation, which lead to differences in performance (e.g., [15,26,30]). Beyond impairing the environment’s perceived realism, the missing cue signals in VR may lead to changes or disruptions in the underlying neural processes in the brain related to spatial memory (e.g. [1]). Furthermore, subjects, especially elderly ones, often have difficulties in performing virtual tasks. Both of these real and virtual world challenges are compounded when developing tools for spatial memory rehabilitation, where ideally subjects should be able to practice in their own homes by themselves and which require much longer sessions.

Augmented Reality (AR) tools have increased in availability and performance in recent years for many different purposes [2,35]). When using Augmented Reality tools, a user views virtual (or “augmented”) objects overlaid on the real world, and this hybrid environment can be viewed via interfaces such as smart-glasses, smartphones and tablets [19]. This enables users to walk around any regular environment, which can be augmented via computational means with targets, landmarks and more. Thus, AR offers us a solution for studying spatial memory with the advantages of both real world and virtual paradigms. AR allows users to naturally move through their environments, while also providing experimenters the flexibility offered by virtual paradigms by having virtual objects and landmarks placed within a real environment. However, as AR technology has only recently become commonly available, to the best of our knowledge no one has yet to perform baseline experiments studying the differences between spatial memory performance in virtual and augmented reality: Is spatial memory similar across conditions? And how does the shift to AR paradigms affect user experience? Do any effects of AR require actual walking or is pointing enough?

As the user experience in AR may appear closer to real world navigation compared to VR ones [19], the use of AR for experimentation might close some of the currently existing gaps between VR and real world tasks in terms of user experience and spatial memory accuracy. Thus, we predict that performance and user experience in AR should be at least equivalent, or superior to, performance in VR, with the added perceptual cues capturing additional relevant behavioral features and moving it closer to natural real-world performance. If true, this would recommend the broader use of AR paradigms for use in spatial memory research and rehabilitation applications.

## Related Work

To our knowledge, spatial memory performance has not been directly compared between AR and VR using matched tasks. However, the relationship between AR and VR has been explored for many other realms, with emphasis on education and training. These include testing educational applications, such as teaching about recycling [39], the water cycle [38], multiculturalism [39], forensic medicine [37] and English as a second language [40]. These studies all found that the use of AR was equivalent to, or better than, VR for the educational tasks tested.

While AR and VR have not been directly compared for spatial memory, two other types of spatial memory comparisons involving VR are relevant to our question. First, performance in VR has been compared extensively to real-world performance, demonstrating the potential for similar levels of accuracy and for transfer between training in one to the other [21,30,33]. From the neuroscience perspective, the extent to which the neural signals underlying behavioral performance are similar between virtual and real-world environments it is currently debated, with some studies showing that signals are maintained while others finding significant differences [1,3,34].

Secondly, in recent years as immersive VR setups that enable physical movement have become available. These include HMD setups with either natural walking in small safe environments, or on an omni-directional treadmill or simply stationary while allowing naturalistic head movements. The use of these setups has been compared to traditional screen-based desktop VR. This line of work has shown general equivalence between the methods, with various advantages to walking over stationary conditions [23,26,27,28,30]. From the neuroscience perspective, it has been suggested that naturalistic head movements may sufficient to elicit natural spatial signals in VR that might be missing in fully virtual paradigms [2]. Note however that these types of walking with an HMD, especially on a treadmill, may still hold considerable subjective difference then realistic real-world walking (e.g. [27,29]) and we would thus expect performance in AR to be between performance in immersive VR to performance in real-world paradigms.

To understand the landscape of research on this topic, we searched for work testing spatial memory performance in AR. We found one previous study that tested the use of AR for spatial memory research. Juan et al. [36] devised “ARSM,” an augmented-reality spatial memory test for children, and showed that it elicits performance patterns that correlate with those seen with more traditional measures. However, this study had limitations because the task used physical fixed points for augmented objects (via QR codes), and did not compare subject’s performance to a virtual version of the same task. Similarly, Khademi [11] and Hondori [10] used AR for spatial-motor rehabilitation, but did not include a memory element or a direct comparison to VR.

In summary, existing literature suggests that while VR is an effective tool for spatial memory research and training, it still has a gap from real world performance. This gap is somewhat mitigated by utilizing immersive VR, but not completely. Thus, performance can be described as VR ≤ immersive-VR ≤ Real world. Given AR’s demonstrated advantages compared to VR, we suggest that AR-based experimental paradigms may be able to narrow this realism gap even further, allowing us to perform experiments that elicit a level of performance between immersive-VR and in the real world.

## Methods

### Paradigm

We developed AR and VR paradigms that adapted the “Treasure Hunt” spatial memory task, previously used in [17,18,20,31]. Treasure Hunt is an object–location associative memory task in which subjects are asked to remember the locations of various hidden objects in a virtual environment. Whereas the previous studies had subjects perform Treasure Hunt in a tropical beach environment rendered only in virtual reality, here we asked subjects to perform the same treasure hunt task in a conference room, in both virtual reality and augmented reality implementations (Fig 1).

**Figure 1.**
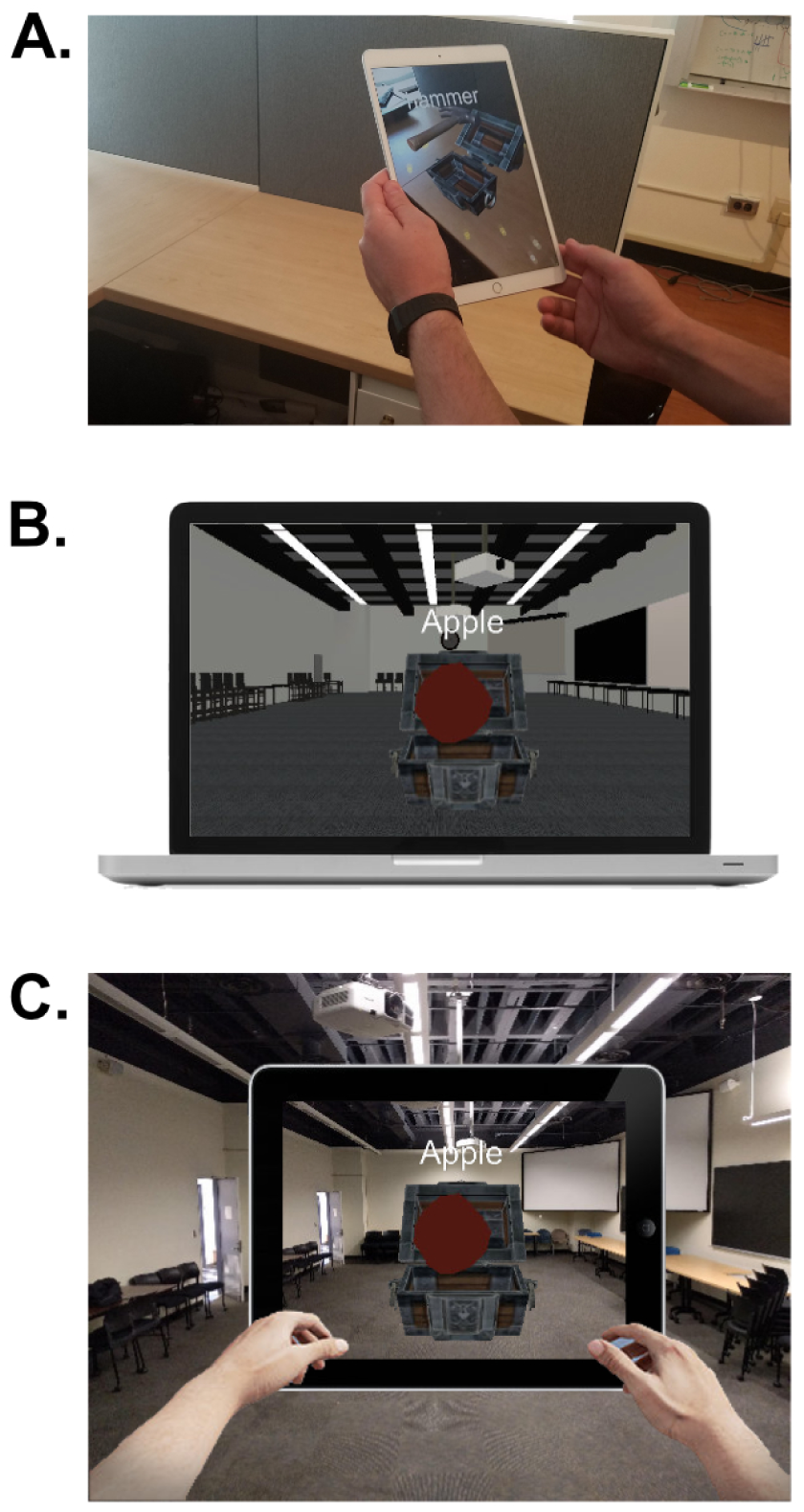
VR and AR. (A) Demonstrating the principle of AR. Note the virtual treasure chest and hammer on the tablet screen overlayed on the desk, while the desk itself is empty. (B) Our desktop VR version of the environment. (C) Our AR version of the task.

In each trial of Treasure Hunt, subjects first perform an encoding phase (Fig 2, Columns 1-2), in which they navigate to a series of treasure chests, each of which is positioned at a random spatial location. When the subject reaches a chest, it opens, revealing an object whose location they are asked to remember. The subject then walks to the next chest. After a series of these learning events, a short distractor phase begins. Here a virtual/augmented rabbit was generated at a random location within the environment and subjects are instructed to follow it. Following the rabbit serves both to distract subjects, forcing them to encode the paired associations, and also to move the subjects away from the location of the third cue (to avoid a scenario where they are cued to recall an object in the location they were currently positioned.) Next, during the retrieval phase (Fig 2, Column 3), patients are shown the name and image of each object and asked to respond by indicating the location where that object was encountered. After recalling the locations of all of the trial’s objects, they receive feedback on their response accuracy. Finally, in the feedback phase (Fig 2, Column 4) the subject views every object’s correct location as well as their response location for each object, with lines linking the two. Here subjects receive points based on their response accuracy and speed in the main spatial-memory task and on their performance on the distractor task. Note that these points were not used in our analysis, as their purpose was mainly to make subjects pay more attention to the distractor task, and finish the overall task in a reasonable amount of time.

**Figure 2.**
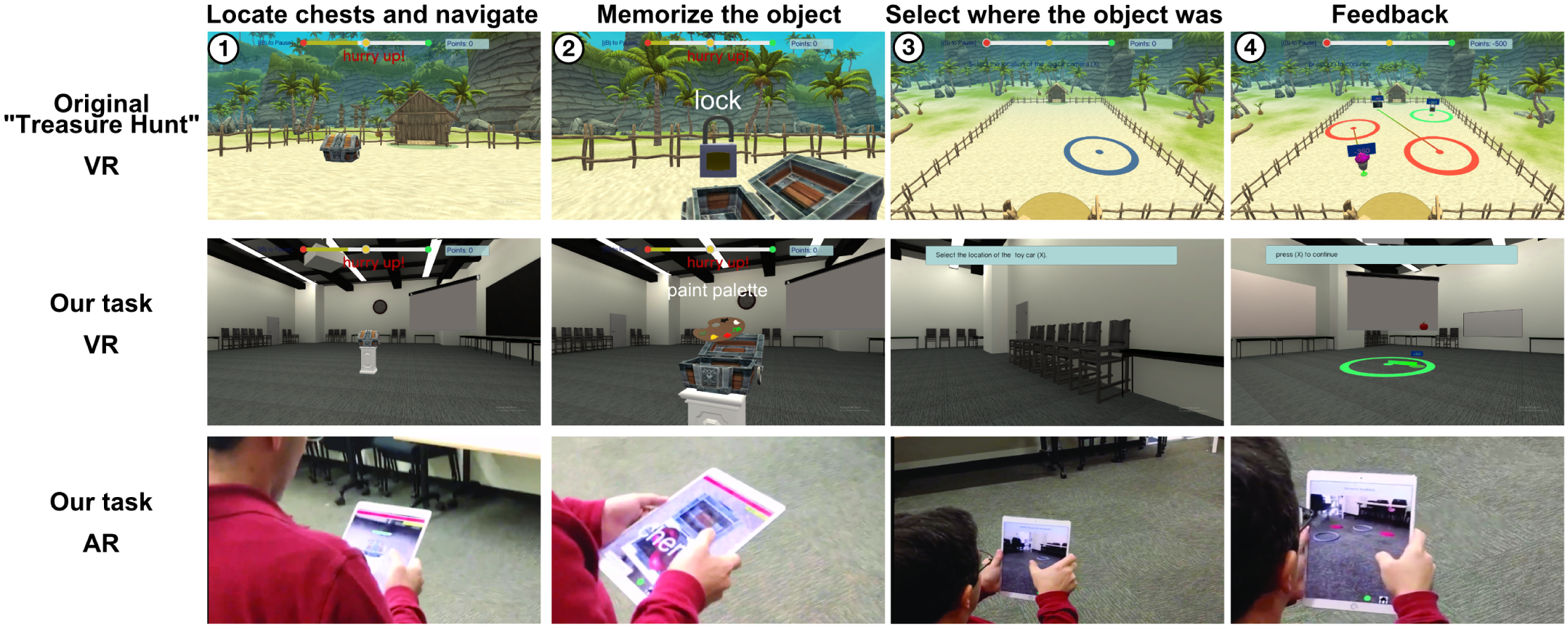
Paradigm. The top row’s screenshots are from the “Treasure Hunt” task on which we based on paradigm. The middle row is from our adaptation of this task for the one used here in virtual reality, and the bottom row from the Augmented Reality version. In each of these versions, subjects perform a series of trials. In each trial they first perform an encoding stage in which they (1) locate 2-4 chests and navigate to them, (2) and then memorize the objects hidden in them. This is followed by a recall stage where subjects are cued to (3) mark the location of specific objects and by a feedback stage(4) in which the true location and the subjects selections are revealed.

Subjects performed 20 trials of the task in each AR setting. Each trial probed either 2 or 3 target chests and 1-2 empty chests. Thus each subject viewed a total of ∼ 50 spatial memory targets for each condition. The overall experiment took subjects ∼ 90–120 minutes, including the time needed to complete a questionnaire and to walk between the rooms where the AR and VR conditions took place. Approximately half of the time was spent on the virtual conditions and half on the augmented conditions. Subjects used a handheld tablet to interact with the environment in the AR setting and a standard desktop screen and keyboard for the VR condition.

To test whether any potential effects stem from physical walking vs. the use of AR in general (i.e are head movements enough or do we need full body walking?), subjects performed both AR and VR task in 2 different conditions, “walking” and “pointing”. In the Walking conditions, we directly adapted the existing “Treasure Hunt” task to AR, using a matched large-room environment in VR and AR (Fig 1 middle and bottom row; Fig 2). In the AR condition, subjects had to physically walk around the room, while in the VR condition they used a joystick to virtually move their avatar. In the Pointing condition, subjects performed the same task but they viewed the environment from a fixed location. Here they pointed at, rather than walked to the targets. In the AR condition they sat in a chair and moved a tablet to look at augmented stimuli across the environment, clicking on the view of the environment through the tablet surface to indicate their responses. In the VR condition they rotated the virtual viewpoint using the keyboard and selected target locations on the screen with the mouse. As the walking conditions required physically/virtually moving through the environment they were inherently slower than the pointing conditions.

### Implementation

#### The VR task

The VR version of Treasure Hunt was developed for Windows using Unity3D game engine (Unity Systems, USA). We replicated the real-world testing environment used in the AR version of the task using 3D modeling software, such as Maya (Autodesk Inc., USA). During the process of creating a virtual environment that served as a replica of the AR testing environment, we were careful to preserve the dimensions of the room as well as the arrangement of different objects like chairs and tables along the peripheral walls of the environment.

#### The AR task

The AR version of the task was developed for iPad using Unity3D and ARKit (Apple Inc., USA), the latter of which is Apple’s library for allowing development of augmented reality applications for iOS devices. ARKit uses a technique called “visual-inertial odometry”, which combines motion sensing information from the accelerometer of the iOS device with computer vision analysis of the scene visible to the back-facing camera. It recognizes notable features in the scene image, tracks differences in the positions of those features across video frames, and compares that information with accelerometer data. Using that information, it is able to track the position of the subject holding the iPad accurately within the augmented reality coordinate space.

For the purpose of creating a testing environment for our AR experiment that would align augmented and real-world landmarks in a consistent fashion across sessions, we utilized a feature of ARKit called “world map,” which saves the tracked features of a scene in the form of a point cloud. When this world map is loaded at the beginning of each experiment, ARKit attempts to match the point cloud data with the features currently being tracked. We note that while the current registration process is relatively smooth (typically 10-30s per experiment) it still leaves significant room for improvement and automation in environments under different lighting conditions and outdoors, which is why we chose an indoor environment for this experiment.

### Participants

24 healthy subjects performed the experiment. Due to a technical issue, the logs for the AR condition for four subjects were not usable. For these subjects, we still used their full VR results and their questionnaire answers. The experiments were approved by Columbia University’s institutional review board (AAAR5000) and all subjects gave their informed consent.

### Statistics

#### Corrected error distance

Our main measure of performance was how accurately the subject remembered the location of each object. To measure this, we computed the corrected error distance between the selected location and the target location. To compute this measure, we first calculated the raw distance for each target, which is the Euclidean distance between the coordinate of the location the subject selected for their response and the actual target object’s coordinate. We then corrected this distance metric, which was important because different target locations can be biased to lead to different potential errors (e.g., if the target is in the center, the maximum error distance can be at most half of the diagonal, while for a target in the corner of the environment the maximum distance can be the full diagonal length of the environment, See Fig 3. For more information on this method see [20]). This correction also controls for potential differences in perceived size and scaling between virtual units in the virtual and augmented environment. Thus, we then corrected the raw distance error by comparing it to the distances between 100,000 points randomly generated inside the environment and each target and assigning the percentile as the corrected error distance.

**Figure 3.**
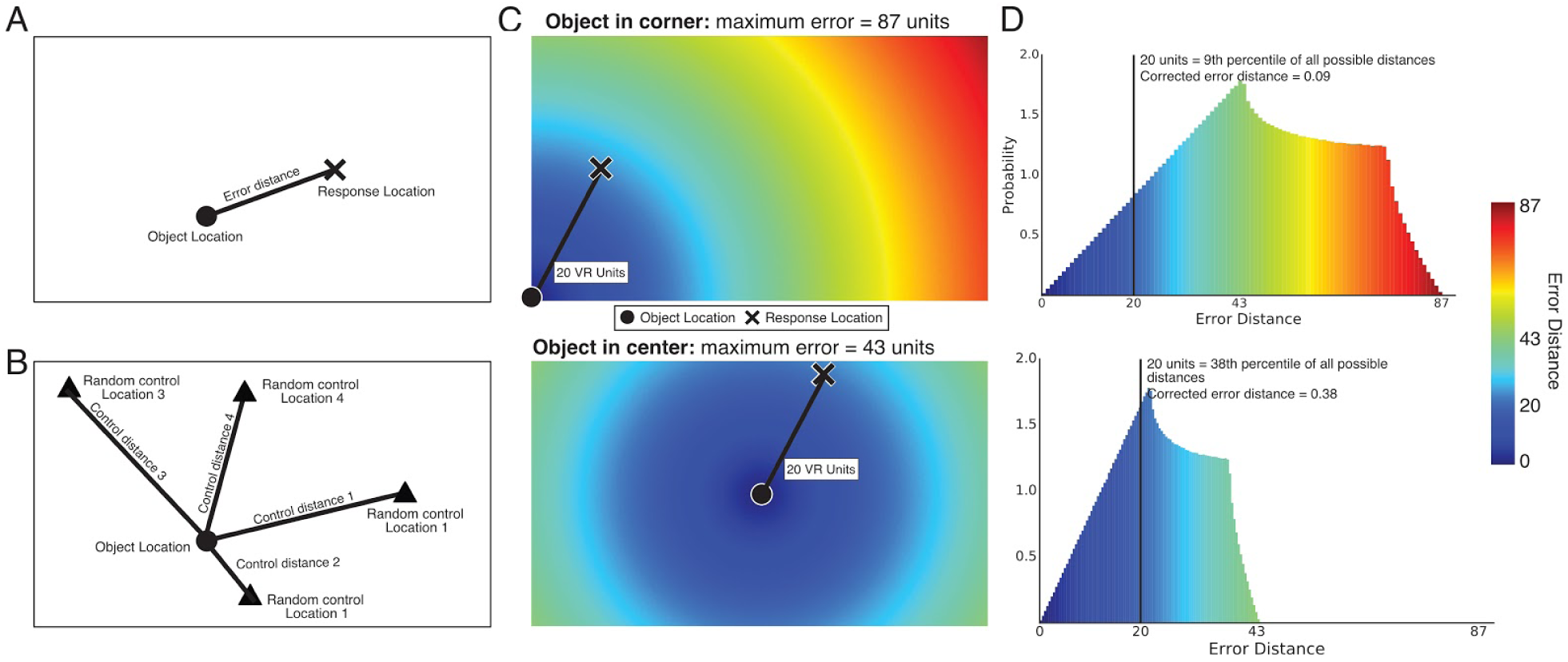
Corrected Error Distance. (A) We first calculated the raw euclidean distance between the object location (circle) and the response location (X) for each object. (B) We then randomly selected 10,000 control locations (triangles) and calculated the distances from each one to the target location (circles). (C) Demonstrates the importance of this correction - when the target is in a corner (top) the range of potential distance errors is much greater than when it is in the center (bottom). (D) We then took the relative percentile within this control distribution as the corrected error distance, which was our main performance measure.

#### Statistical comparisons

To statistically compare the subjects’ memory performance **between conditions**, we calculated the mean corrected error distance per subject in each condition, and then performed a paired signed-rank test.

In order to test the **overall significance of a subject’s performance** in a given condition, we generated two values for each trial - the subject’s raw distance score and a trial-specific chance value generated by taking the distance at 50% in the permutation measure.. We then tested the relationship between the pairs of selected distances and the chance ones using a signed-rank test. We refer to this procedure as “spatial memory scoring” or simply “SMS test.”

This metric and these analysis will enable us to directly test in the next section whether AR and VR lead to significant spatial memory performance for each subject, and to compare their relative performance across the four conditions.

## Results

In order to understand spatial memory in AR, subjects performed our version of the “Treasure Hunt” spatial memory task in each of four conditions: (in a pseudo-random order). Each task was performed in both AR and VR settings and in both freely walking and pointing modalities (fully crossed for four possible combinations). We assessed subject performance in all four conditions by measuring spatial memory accuracy and for user experience. Subjects also performed a standard questionnaire that assessed their spatial abilities (The Santa Barbara Sense of Direction scale, SBSoD [8]).

### Performance when walking in AR and VR

We first tested whether subjects could perform the task well by assessing whether their memory accuracy was above chance. In the AR condition the mean memory accuracy was 0.09±0.01 (normalized units). We found that the subjects all were able to perform the task significantly above chance (all p’s<10^−7^, SMS test). This demonstrated that subjects were consistently able to respond at locations relatively close to the actual memory target position.

We then measured subject performance while they performed the task in the VR walking condition. We found that here too subjects were able to perform the task significantly above chance (mean accuracy = 0.16±0.01, all p’s<0.03, SMS test). We also compared their performance to a wider baseline of data from the standard implementation of the “Treasure Hunt” task from ([16], see first row of Fig 1) and found that the results were in line with this baseline (p=0.65, unpaired two-tailed t-test). This demonstrates the validity and fit of our task for successful testing spatial memory. It also demonstrates that behavioral performance was in line with previous results despite changes in the task here vs. the baseline task (e.g. indoor vs. outdoor environment).

### Comparing AR and VR

We next compared the memory accuracy of subjects who performed the walking version of the task in AR versus VR settings. Here we found that subjects were significantly more accurate in AR then in VR (mean memory accuracy was 0.09±0.01 and 0.16±0.01 respectively, p<0.001) (Fig 4).

**Fig 4.**
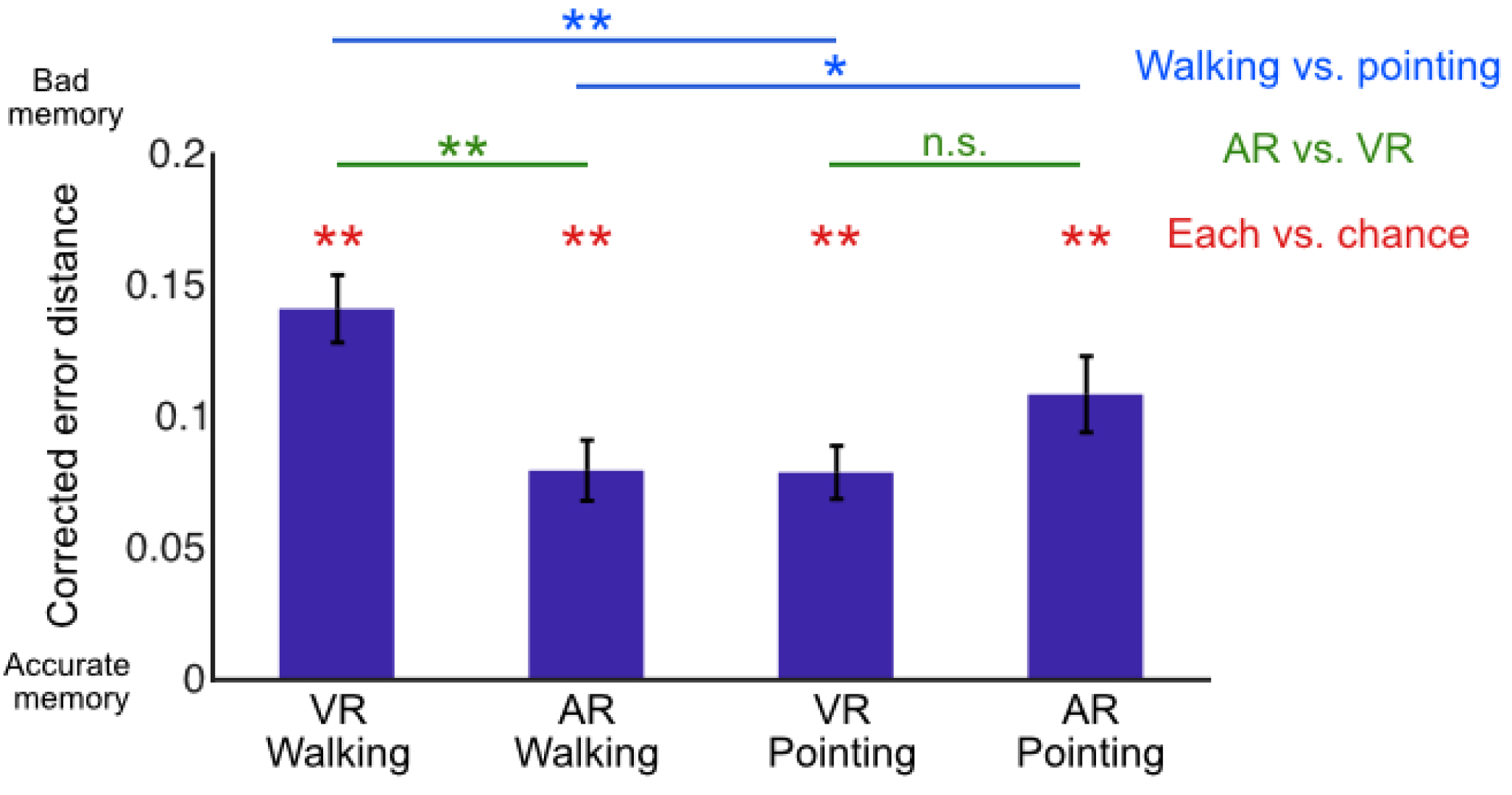
Spatial Memory accuracy. Subjects showed significant spatial memory in all 4 conditions. When comparing AR to VR, we found a significant advantage when walking, but not when pointing. When comparing walking to pointing within each condition we found a mixed result, with better performance in AR for walking and in VR for pointing.

We then compared subjects’ subjective experience between the two conditions (Fig. 5). Subjects subjectively reported that the AR version was easier (Means = 2.8±0.3, 4.3±0.2 respectively, p<5.6 × 10-6), more enjoyable (Means = 3.7±0.3, 2.8±0.3 respectively, p<0.01) and more immersive (Means = 4.1±0.2, 3.3±0.3 respectively, p<0.05) then the VR task. This was joined by the subjects subjective descriptions - “Overall, the mobile AR was fun and immersive” S53 “When I feel disconnected from my body, I had difficulty to estimate my location accurately. “S45 “to sense the space in VR is much harder.” S70

**Figure 5.**
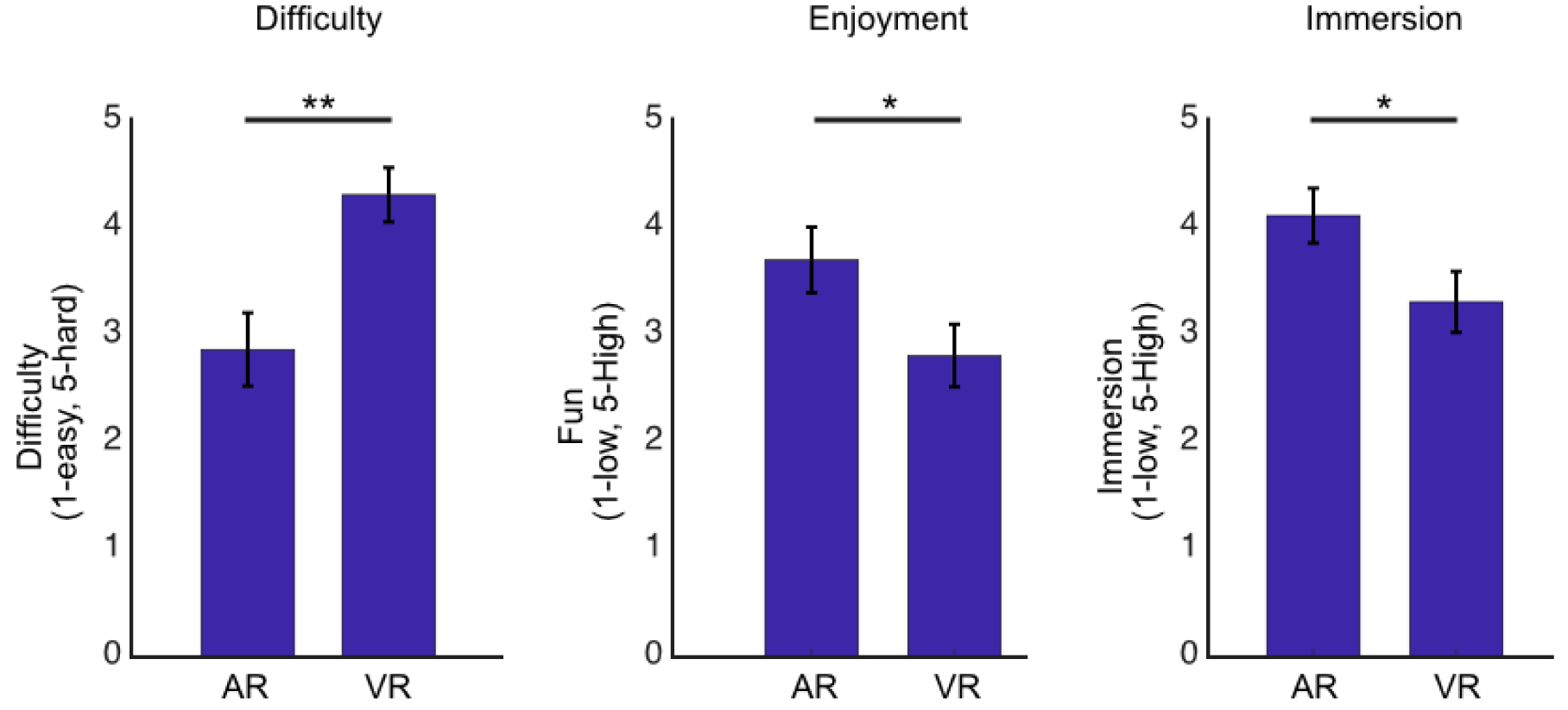
AR vs. VR. Subjects subjectively reported that AR was significantly easier, more fun and more immersive than VR.

Demonstrating the measured memory performance is elevated in AR and that subjects subjectively prefer AR over VR in terms of difficulty, immersion and enjoyment is the main result of this project, indicating that assessing spatial memory in AR is at least as valid, if not significantly better, then VR. Whereas previous VR memory paradigms had been criticized because memory performance was lower compared to real-world spatial memory [30], our results show that AR helps address this gap.

### Performance when pointing

We also tested subjects’ performance when pointing from a stationary position instead of walking, in both AR and VR. We found that all subjects’ performances were significantly above chance when pointing in both AR (all p<10^−15^, SMS test) and VR (all p’s<10^−4^, SMS test).

We compared the subjects spatial memory accuracy in pointing between AR and VR, and found no significant difference,but a trend towards VR (p=0.07). In terms of user experience, there were no significant differences between AR and VR in terms of difficulty (p=0.9), enjoyment (p=0.25) or immersion (p=0.15). Notably, subject’s performance and opinions varied widely, with some showing a strong preference for AR and some for VR. This was reflected by the user comments, for example: “Stationary VR was not as easy as Stationary AR” (S47), “the virtual version is much more easier then AR” (S73).

The lack of a clear difference between AR and VR when pointing amy suggest that the ability to physically move around and reach the target may diminish the effect we reported above for walking.

### Performance when walking vs. pointing

When comparing memory performance between the walking vs. pointing conditions, we found that whereas in AR accuracy was better when walking then when pointing (0<0.05, signrank), in VR pointing was more accurate than walking (p<0.01, signrank). Here too subject’s subjective opinions were split, for example:”Being stationary made it more difficult to recall the placement of the objects because it was hard to tell how close or how far away things were” (S57), “felt slightly easier than the mobile version, which was difficult because of directional control” (S46)

However, due to the substantial task differences between responding based on pointing versus walking, it is hard to disentangle whether this difference comes from a true difference in spatial memory, or from a perceptual and interface difference related to the accuracy of pointing at a remote spot on a 2d screen (whether desktop or AR-tablet) vs. physically walking there in 3D, especially in the real world which gives better depth perception. Nonetheless, this pattern of results points at the potential importance of movement for spatial memory, continuing the previous finding, possibly especially for depth perception (e.g. “Sitting down diminished my sense of depth/closeness of the chests.” (S51)).

### Spatial memory performance and sense of direction

We used the Santa Barbara Sense of Direction (SBSoD) scale questionnaire to assess how each subject perceives their own spatial abilities [8]. This measure has been shown to strongly correlate with many other spatial measures (e.g. [4,8,9,32]). Therefore, we then correlated subjects’ scores on the Santa Barbara questionnaire with their performance in each of the conditions. While we did not find a correlation between SBSoD to VR performance when walking (r=-0.13) or pointing (r=0.07) or the pointing in AR (r=0.06), performance while walking in AR showed a moderate correlation (r=0.4, p<0.05). This result was also consistent with subject’s subjective responses. Several subjects reported that AR felt closer to natural behavior. For example: “I felt like I was doing something totally different when actually walking, this just felt natural” (S62), “In VR I felt like my body was not connected to my movements and I was totally disconnected” (S45).

Despite these results, note that there was no correlation between accurate performance in walking AR to feelings of difficulty, enjoyment and immersion (all correlations p>0.49). This suggests that walking in AR may be better at capturing spatial skills and performance than the other conditions, possibly due to being more close to naturalistic real world spatial behavior. It also demonstrates that AR’s improved accuracy and link to self-perception of spatial skills are not linked to subjective difficulty, enjoyment and immersion in AR, showing higher spatial accuracy also when subjects lack positive feelings about the task.

## Discussion and future work

Our main novel finding was that subjects show better spatial memory performance while performing a task while walking in AR, compared with performing a comparable task in VR. Further, subjects found the AR version of the task to be significantly easier, more fun, and more immersive. We additionally found that these AR versus VR differences were diminished when subjects indicated their responses while pointing instead of walking, suggesting that the naturalistic physical movement may play an important role in spatial memory.

It is well established in the literature that spatial cognition performance is worse in VR environments compared to the real world (“VR<=Real”), with this gap remaining even with immersive VR. The VR performance gap indicates that some key spatial knowledge is not perceived by subjects with existing VR tools. In line with the Milgram’s definitions of the virtual spectrum [19], our results suggest that AR may narrow the gap from real world performance even further (VR<=immersive-VR<=AR<=Real), bringing us closer to natural performance. By helping to close this performance gap, as well as by bringing fun and immersion to navigational experience, it suggests that AR may have substantial potential for spatial memory research and rehabilitation.

In this context, it is important to note a limitation of our work, which is that we compared spatial memory performance with AR to desktop VR, not to immersive VR, which enables physical walking around in a room or on a treadmill (see Introduction). It is possible that immersive VR would show improved performance, closing the gap with our AR findings. Furthermore, while AR has the advantages of both real world and immersive VR, it is still in its current technological level a compromise between them. Although AR provides much more flexibility than regular real-world environments, it still does not match the flexibility of fully immersive VR as it is continues to rely on the basic layout of the actual physical environment. The naturalistic feeling from using AR tends to break down in complex environments with uneven surfaces where sometimes the accuracy of the positioning of augmented objects can be problematic. In both of these cases, further advances in AR technology will continue to mitigate these differences to a great extent [6,7,12,22]. Thus, though AR tools are still new and evolving and we can expect improved results going forward, even current versions can already be utilized to create experiments that are more naturalistic and better capture human performance.

### Potential for Research

Following earlier advances, AR has the potential of being an extremely powerful tool for psychological and neuroscience research. An important first step, which we contribute to here, is in establishing clear behavioral baselines for performance in AR, to enable better extrapolation and generalization from the much larger existing VR research. Specifically, for the research of spatial memory, one can use current AR tools to test a range of questions in spatial memory research. For example, Will we see differences between familiar and unfamiliar environments? How does memory performance in AR environments vary indoors versus outdoors?

The greater ecological validity of AR can offer especially strong potential when combined with mobile neuroimaging (e.g. mobile fNIRS, mobile EEG) and neuro recording (e.g. the chronically implanted Neuropace [14] or the RC+S devices [13]). This can enable us to create flexible, but highly controlled, paradigms in naturalistic real world settings, which might allow us to identify novel brain signals that have been previously missing from findings obtained from VR-based paradigms [1].

### Potential for Rehabilitation

Current research approaches for spatial memory rehabilitation face similar kinds of challenges as spatial memory research, although often the magnitude of these problems is magnified by the need for the paradigms to be accessible to subjects with memory impairments [12,25]. Existing real-world paradigms are often too cumbersome to run in the clinic, not to mention home, and virtual paradigms have not been successfully adopted. Because AR may be more convenient, intuitive, and enjoyable for individuals with memory impairments, these methods may have special utility for working with these challenged patients—this is a view that has also been previously suggested by others in pointing out the potential of AR for rehabilitation (e.g. [6,11,12,24]). Furthermore, beyond the advantages mentioned above regarding naturalness and flexibility, our findings show also that AR has the advantage of being easier to use. For all of these reasons, we see great potential in future use of AR for spatial memory rehabilitation.

## Conclusion

Our main impact and novelty is in providing a first quantitative measurement of the improvement in spatial memory accuracy that results from performing a single task in both augmented and virtual environments. Our finding that spatial memory encoding in AR was significantly easier, more immersive and more fun as compared to VR, and that performance was significantly more accurate, demonstrates that AR may be an improvement compared to VR for spatial memory research and rehabilitation because it will lead to more natural spatial performance. These findings hold significant potential as a foundation for future spatial memory research and rehabilitation.

## Acknowledgments

This research was supported by NIH Grant MH104606 and the National Science Foundation (JJ) and by NIH F32 MH120990 (SM). We additionally wish to thank our subjects for participating in our experiments, and Shi-Fu Chang and the Columbia University Fu Foundation School of Engineering for giving us access to the experimental space.

